# Lipid metabolism dysfunction following symbiont elimination is linked to altered Kennedy pathway homeostasis

**DOI:** 10.1101/2022.08.23.504685

**Authors:** Geoffrey M. Attardo, Joshua B. Benoit, Veronika Michalkova, Alekhya Kondragunta, Aaron A. Baumann, Brian L. Weiss, Anna Malacrida, Francesca Scolari, Serap Aksoy

## Abstract

Lipid metabolism is critical for insect reproduction, especially for species that invest heavily into early developmental stages of their offspring. The role of symbiotic bacteria during this process is unknown but likely essential, especially in the case of obligate microbes that fulfill key biological functions in the host. Using a combined lipidomics, functional genomics and biochemical strategy, we examined the role of lipid metabolism in the interaction between the viviparous tsetse fly (*Glossina morsitans morsitans*) and its obligate endosymbiotic bacteria (*Wigglesworthia glossinidia*) during tsetse pregnancy. We observed increased *CTP:phosphocholine cytidylyltransferase* (*cct1*) expression during pregnancy. This gene codes for the enzyme that functions as the rate limiting step in phosphatidylcholine biosynthesis in the Kennedy pathway which is critical for stored lipid metabolism and progeny development. Experimental removal of *Wigglesworthia* impaired lipid metabolism via disruption of the Kennedy pathway, yielding obese mothers whose developing progeny ultimately starve. Functional validation via experimental *cct1* suppression revealed a phenotype similar to females lacking obligate *Wigglesworthia* symbionts. These results indicate that, in *Glossina*, symbiont-derived factors, likely B vitamins, are critical for proper function of both lipid biosynthesis and lipolysis. Loss of the symbiosis has a dramatic impact on *Glossina* fecundity, and may be broadly applicable to other insect systems, particularly to species that require symbiotic partners to maximize lipolysis and reproductive output.

## Introduction

Reproduction represents an evolutionarily imperative and metabolically taxing physiological process for all animals (Flatt, 2011; Perrin and Sibly, 1993). This may be especially true for insects, which maintain a high reproductive rate, even though progeny quality declines throughout adulthood (Barreaux et al., 2022; Michalkova et al., 2014a). One such insect is the tsetse fly, *Glossina spp*., which reproduces via a process called ‘adenotrophic viviparity’ (gland fed, live birth (Benoit et al., 2015)). Female tsetse ovulate one egg per gonotrophic cycle and following fertilization and embryogenesis, larvigenesis proceeds within the maternal uterus. *In utero*, larvae are nourished exclusively by milk secretions produced by uniquely adapted maternal accessory glands, called the milk glands. Thus, female tsetse flies supply all the nutrients necessary to support embryonic and larval development of their progeny. Tsetse milk consists of 20-30 mg of carbohydrates, lipids and proteins in an aqueous base (Benoit et al., 2015, 2014; Cmelik et al., 1969; Tobe, 1978). The rapid incorporation of nutrients into milk products during lactation reduces total maternal lipid and protein content by nearly 50% and 25%, respectively (Attardo et al., 2012). Following birth, the milk glands rapidly involute and lipid reserves accumulate to support a subsequent offspring (Attardo et al., 2012; Baumann et al., 2013; Benoit et al., 2018). Tsetse mothers endure this metabolically intense K-selected reproductive strategy continuously through their adulthood, ultimately birthing up to 8-12 larvae per lifetime (Barreaux et al., 2022; Michalkova et al., 2014a).

Tsetse flies have evolved to feed exclusively on vertebrate blood. While blood is nutrient-rich, it lacks in sufficient quantity several of the vitamins and cofactors required to support the metabolically intense process of milk production. To overcome this, tsetse flies have developed intimate, long-term associations with symbiotic bacteria that supplement nutrients missing from host blood. One of these bacteria is the maternally transmitted obligate endosymbiont *Wigglesworthia glossinidia*, which has associated exclusively with tsetse species for 50-80 million years (Chen et al., 1999). As a result, *Wigglesworthia* has experienced massive genome erosion (Akman et al., 2002; Hall et al., 2019). However, the bacterium retains genes required to produce a variety of B vitamins that are largely absent from vertebrate blood (Akman et al., 2002; Bing et al., 2017). Metabolic processing of proline, which circulates in tsetse hemolymph and replaces carbohydrates as the flies’ prominent energy source, is fueled by B vitamins generated by *Wigglesworthia* (Michalkova et al., 2014b). Accordingly, antibiotic-mediated clearance of *Wigglesworthia* results in a dramatic reduction in multiple B vitamin-associated compounds that serve as essential cofactors in fundamental metabolic pathways, including the pentose phosphate pathway, nucleotide biosynthesis, methionine/cysteine metabolism, and amino acid and lipid metabolism. These deficiencies yield adult female flies incapable of milk production which consequently results in abortion of early instar larvae (Benoit et al., 2017; Bing et al., 2017; Nogge, 1976). However, the mechanistic basis for this phenotype remains unclear.

Here we examined the interplay among obligate symbionts, lipid metabolism, and reproductive output of tsetse flies. Maternal transfer of nutrients to developing offspring is an energetically demanding process for all insects. However, tsetse flies, similar to other viviparous systems, invest even more into their progeny by providing resources for embryonic and larval development. This study demonstrates that critical symbiont-derived factors promote phospholipid synthesis, facilitating rapid lipolysis of stored lipid reserves in order to nourish the developing intrauterine progeny. Other insects that rely on symbiont-derived B vitamins, including bed bugs and lice (Benoit et al., 2016; Boyd et al., 2017; Hosokawa et al., 2010; Kirkness et al., 2010), are likely to experience impaired lipolysis and reduced fecundity in the absences of their symbiotic partner(s), as the molecular mechanisms underlying this process are not unique to *Glossina* (Thomas et al., 2020). Furthermore, similarities in live birth across animal systems indicate that micronutrient deprivation is likely to have significant impacts on metabolic processes with similar negative consequences that extend from invertebrates to vertebrates (Fouks et al., 2022).

## Results

### Aposymbiotic flies are obese and fail to reproduce

Antibiotic-treated, endosymbiont-free (aposymbiotic) tsetse flies failed to reproduce successfully, as previously shown (Benoit et al., 2017; Michalkova et al., 2014b; Nogge, 1976). When early instar larvae from aposymbiotic mothers were removed before abortion and compared to control larvae, their lipid levels were 25-30% lower, indicating substantially reduced lipid transfer from mother to larvae in the aposymbiotic state (Fig. 1). Aposymbiotic females show higher total lipid levels, with an average of 5.66 ± 0.44 lipids per mg dry mass relative to 4.85 ± 0.34 in symbiotic females. This phenotype was associated with increased lipid droplet (LD) size in the fat body (FB), suggesting defect(s) in lipolysis. Additionally, Nile blue staining of FB tissue revealed a reduction in charged phospholipids relative to neutral storage lipid species, including tri- and di-glycerides, in aposymbiotic flies relative to control flies (Fig. 1). These initial studies suggest a major dysfunction in lipid metabolism following symbiont clearance.

**Figure 1:**
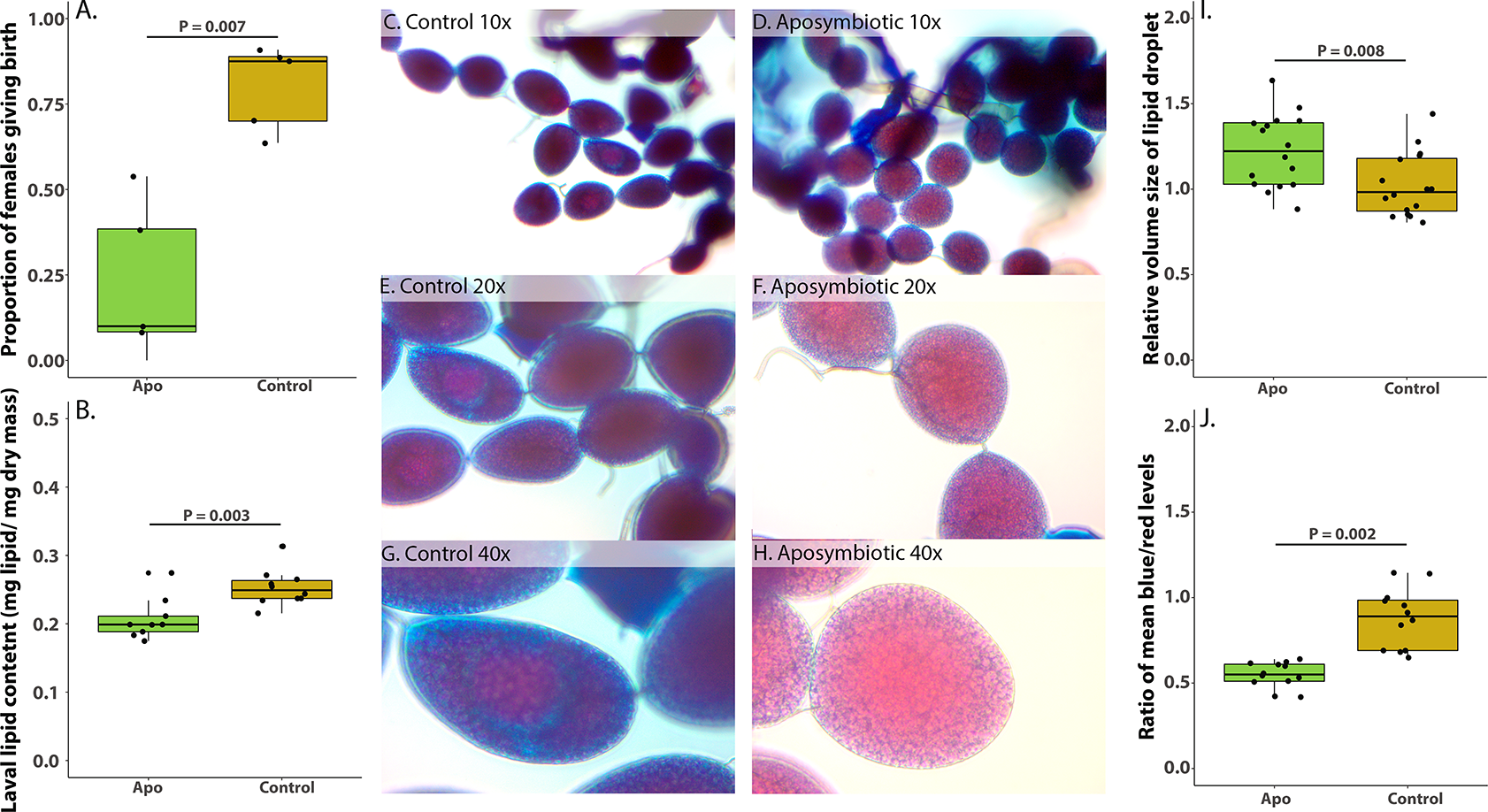
Reproduction and lipid metabolism is dysfunctional in aposymbiotic flies. Proportion of females giving birth (A), N = 4, and lipid content in the surviving larvae (B) are significantly reduced in aposymbiotic flies (t-test, P-value 0.007 and 0.0003, respectively). N = 8-10. C-H) Nile Blue staining reveals significant differences in the neutral and charged lipid composition in aposymbiotic and control flies. I) Volume of the fat body is increased significantly in aposymbiotic flies (t-test, P-value 0.008). N = 12-14. J) Image quantification reveals significantly higher charged lipids in the control compared to the aposymbiotic flies (t-test, P-value 0.002). N = 12-14.

### Lipidomics reveals massive shifts in metabolic lipid processing

To understand the biochemical landscape facilitating the accumulation of triacylglycerols in endosymbiont-free flies, we generated aposymbiotic females via tetracycline application, and analyzed pooled FB and milk gland (MG) tissues, comparing aposymbiotic and untreated (control), age-matched females. Analysis of the differential biochemical composition revealed substantial alterations in the composition of multiple biochemical categories in aposymbiotic flies, including B vitamins, creatine/creatinine, amino acids, diacylglycerols (DAG), phospholipids, sphingolipids, lysolipids and polyunsaturated fatty acids (Suppl. Figs. 1-6). These changes yield an overall composition difference in the lipid profiles between control and aposymbiotic flies (Suppl. Fig. 7, ANOSIM, R = 0.879, P = 1.03E-06).

Many of the observed changes are via either direct or indirect effects of dysregulation of phospholipid (PL) biosynthesis via the Kennedy pathway (Fig. 2). Notably, aposymbiotic flies were deficient in key precursors to PL synthesis including choline, phosphocholine, and the majority of the detected DAGs, and phosphatidylcholines (PCs). Many DAG variants are present in adult female tsetse, and they differ in acyl chain lengths and desaturation state. The most abundant DAGs in control flies contain 16/18 carbon unsaturated or mono-unsaturated side chains (Fig. 2 and Suppl. Fig. 2), while aposymbiotic flies show a broad reduction in abundance of these DAGs. Among phosphatidylcholines, aposymbiotic flies generally lack moieties containing 16/18 carbon unsaturated or mono-unsaturated side chains, while those containing 18/20 carbon polyunsaturated fatty acids are abundant (Suppl. Fig. 1). This may reflect the observed overabundance of polyunsaturated free fatty acids in aposymbiotic flies (Suppl. Fig. 4). The substantial deficiencies in the precursor molecules required for phosphatidylcholine production suggest that *Wigglesworthia*-derived factors either directly or indirectly facilitate the production of phospholipid precursors.

**Figure 2:**
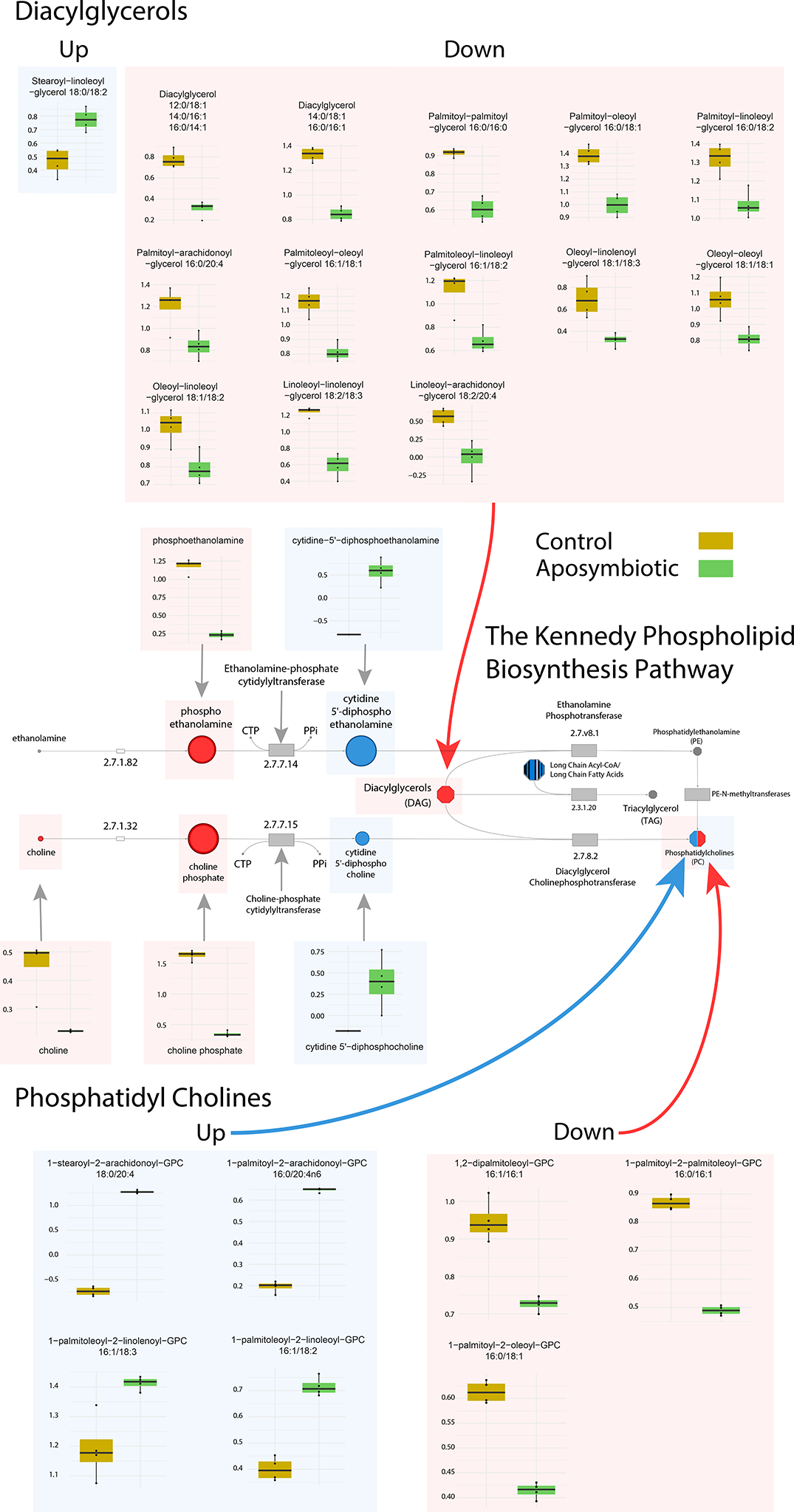
Lipidomics reveals a significant shift in lipid moetities following symbiont removal. Compounds showing a significantly different fold change in their abundance in control (yellow) and aposymbiotic (green) flies are shown. Red denotes increase in compounds of the Kennedy pathway in aposymbiotic flies, while blue indicates lipids showing higher fold change in control individuals. Specifically, there is a major dysfunction in the Kennedy pathway for phosphatidylcholine synthesis. N = 5, * indicates significance based on a t-test.

Aposymbiotic flies also demonstrate a profound deficiency in creatine and creatinine (Suppl. Fig. 6). Creatine/creatinine biosynthesis and the urea cycle are dependent on folate, which is provided to tsetse flies by *Wigglesworthia* (Snyder and Rio, 2015). Components of the urea cycle are also reduced in abundance in aposymbiotic flies. The urea cycle is also dependent on folate for proper function (Brosnan and Brosnan, 2010; Fatterpaker et al., 1951). Creatine/creatinine are key regulators of lipid homeostasis for adrenergic and diet-induced thermogenesis in brown and beige adipose tissue in vertebrates. Induced deficiencies in in creatine/creatinine in mice results in diet induced obesity, which can be rescued by dietary creatine supplementation (Kazak et al., 2017). The mechanism behind how these compounds regulate lipolysis remains to be elucidated but may also be involved in maintenance of lipid homeostasis throughout the reproductive cycle.

### Choline-phosphate cytidylyltransferase 1 represents a critical enzyme during tsetse fly reproductive cycles

Prior RNA-seq analyses of gene expression during tsetse pregnancy and milk production revealed a suite of highly-expressed milk protein-encoding genes (Attardo et al., 2019; Benoit et al., 2014; International Glossina Genome Initiative, 2014). Notably, transcripts for the enzyme choline-phosphate cytidylyltransferase 1 (*cct1*) were upregulated in a similar manner to milk proteins in pregnant flies (Fig. 3). This enzyme participates in phosphatidylcholine synthesis. Phosphatidyl cholines are critical for regulation of LD size and formation to facilitate lipolysis during energy-intensive processes (Krahmer et al., 2011). The expression of *cct1* increases during milk gland involution following parturition (Fig. 3). Tissue-specific expression analysis confirmed *cct1* transcripts were abundant in the FB and MG. In support of this finding, *in situ* hybridization revealed *cct1* expression in both of these tissues (Fig. 3C). These data suggest that *cct1* is critical for lipid metabolism during tsetse pregnancy, likely functioning in the FB and MG, to break down lipids critical as energy for milk production, or necessary to produce milk LDs.

**Figure 3:**
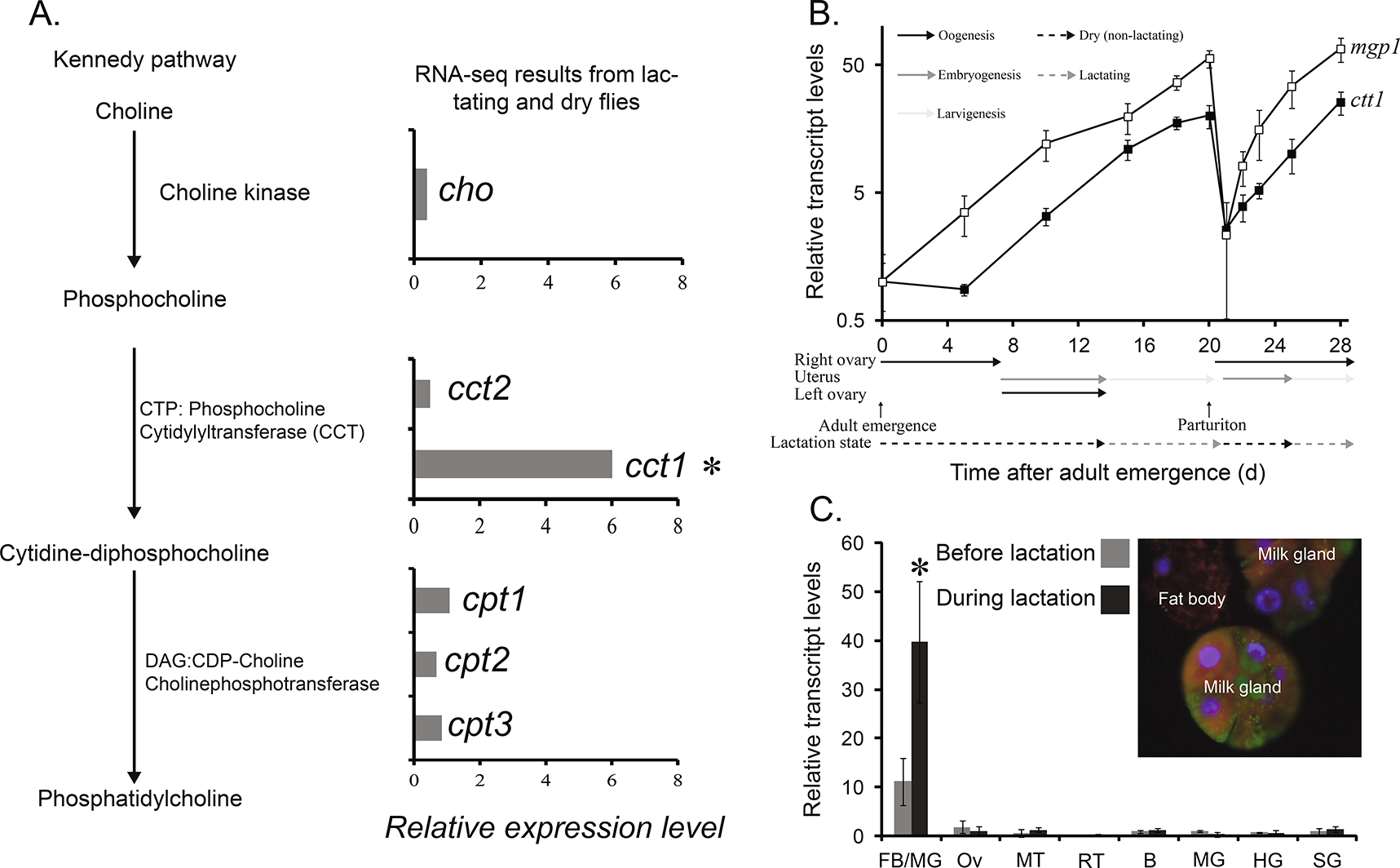
Increased expression of *choline-phosphate cytidylyltransferase 1 (cct1)* is associated with tsetse fly pregnancy. A) Previous RNA-seq studies (Attardo et al., 2019; Benoit et al., 2014) reveal that *cct1* is expressed in pregnant/lactating flies. B) qPCR indicated that the expression pattern for *cct1* is correlated with the pregnancy cycle, similar to a milk protein critical to feed the developing larvae (*milk gland protein 1, mgp1*). N = 4. C) Expression of *cct1* is localized in both the milk glands and fat body, suggesting critical roles in lipid breakdown and milk production. N = 4, *, denotes significance based on a Kal’s proportion test or t-test.

### Suppression of cct1 phenocopies the aposymbiotic state

To confirm if impaired *cct1* function and altered phospholipid metabolism perturbs lipid metabolism during viviparity, we suppressed *cct1* expression using RNA interference (RNAi) (Fig. 4). Similar to aposymbiotic flies, *cct1* knockdown flies produce few progeny (Fig. 4). We noted a massive decrease in levels of phospholipids and PC in developing larvae, indicating that *cct1* is necessary to ensure sufficient lipid transfer to the developing larva. This decrease likely accounts for the 40-45% reduction in total lipids measured from larvae deposited from *cct1* knockdown flies (0.20 ± 0.03 mg lipid/ mg dry mass for *cct1* RNAi vs 0.29 ± 0.04 for control). In aposymbiotic females, *cct1* expression increases substantially, perhaps as a compensatory response to reduced phospholipid levels, which are required for lipid transfer during pregnancy.

**Figure 4:**
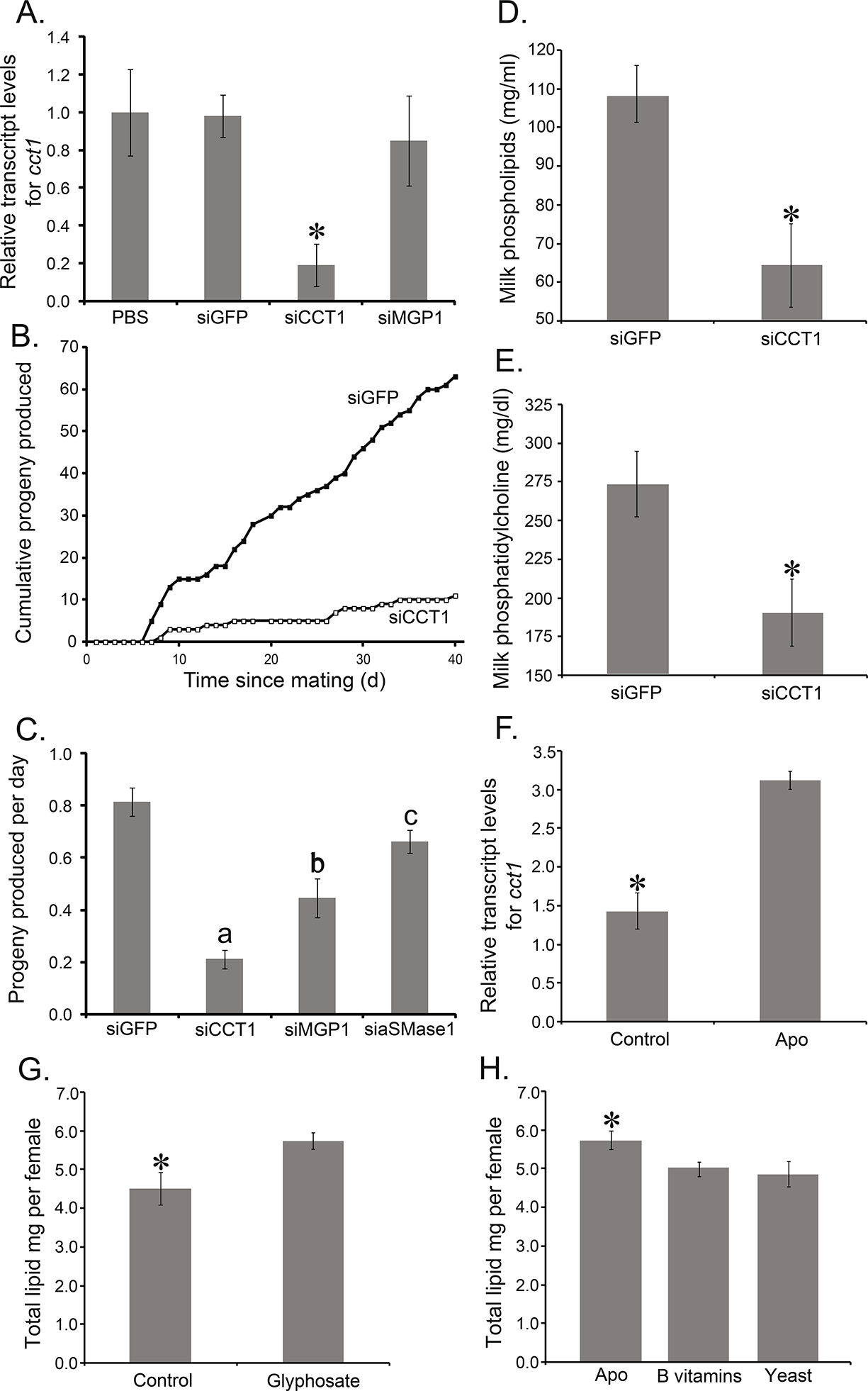
Suppression of *cct1* produces aposymbiotic phenotypes of altered lipid metabolism during pregnancy. A) RNA interference reduces the expression of *cct1*. N = 5, *, denotes significance based on a t-test. *Mgp1, milk gland protein 1.* B-C) Total progeny production and progeny generated per day are reduced following suppression of *cct1*. N = 3 groups of 10. Different letters denote significance based on an ANOVA. *Acid smase1, asmase1*. D-E) Milk phospholipids and phosphatidylcholine are reduced when *cct1* is reduced. N = 12. *, denotes significance based on a t-test. F) Aposymbiotic flies show increased expression of *cct1* as a potential compensatory mechanism to allow for lipolysis during pregnancy. N = 10. *, denotes significance based on a t-test. G) Glyphosate treatment increased lipid levels within flies. N = 8. *, denotes significance based on a t-test. H) Blood supplementation with B vitamins or yeast extract reduced obesity in flies. N = 10. *, denotes significance based on ANOVA.

Tetracycline treatment impairs mitochondrial function (Moullan et al., 2015), which could represent an off-target outcome that significantly impacts tsetse lipid metabolism. As such, we treated flies with glyphosate, which specifically inhibits *Wigglesworthia* folate production (Rio et al., 2019). This yielded flies that presented obesity measures similar to their aposymbiotic counterparts (Fig. 4G). Lastly, aposymbiotic flies were fed on blood with a cocktail of B vitamins and yeast extract, which reduced the lipid levels within the flies, providing a direct link between B vitamins and obesity levels (Fig. 4H). This finding confirmed that *Wigglesworthia*-derived B vitamins acts as an essential factor for tsetse lipid homeostasis.

## Discussion

This study defines a novel feature of host-symbiont dynamics in tsetse showing dysfunctional PL metabolism as a resultant phenotype of symbiont elimination (Fig. 5). Tsetse fly pregnancy requires a rapid transfer of substantial nutritional resources from mother to offspring. A key mechanism in this process is the production of PC by the Kennedy biosynthesis pathway. The *cct1* gene encodes a rate-limiting enzyme in this pathway that prepares the choline headgroup for attachment to cytidyl triphosphate to generate CDP-choline. CDP-choline is then combined with diacylglycerol to produce the final phosphatidyl choline product. The observed increase in *cct1* transcripts during pregnancy suggests that this enzyme/pathway is critical to provision of PC for LD production/function for lipolysis in the FB and for incorporation into products secreted from the MG (Benoit et al., 2015; International Glossina Genome Initiative, 2014). Production of PC is required to coat the outer surface of LDs produced by the endoplasmic reticulum, and is thus critical for lipid metabolism (Olzmann and Carvalho, 2019). The amphipathic nature of PC prevents aberrant LD fusion and provides the biochemical conditions to recruit LD-associated proteins required for lipolysis (Pol et al., 2014).

**Figure 5:**
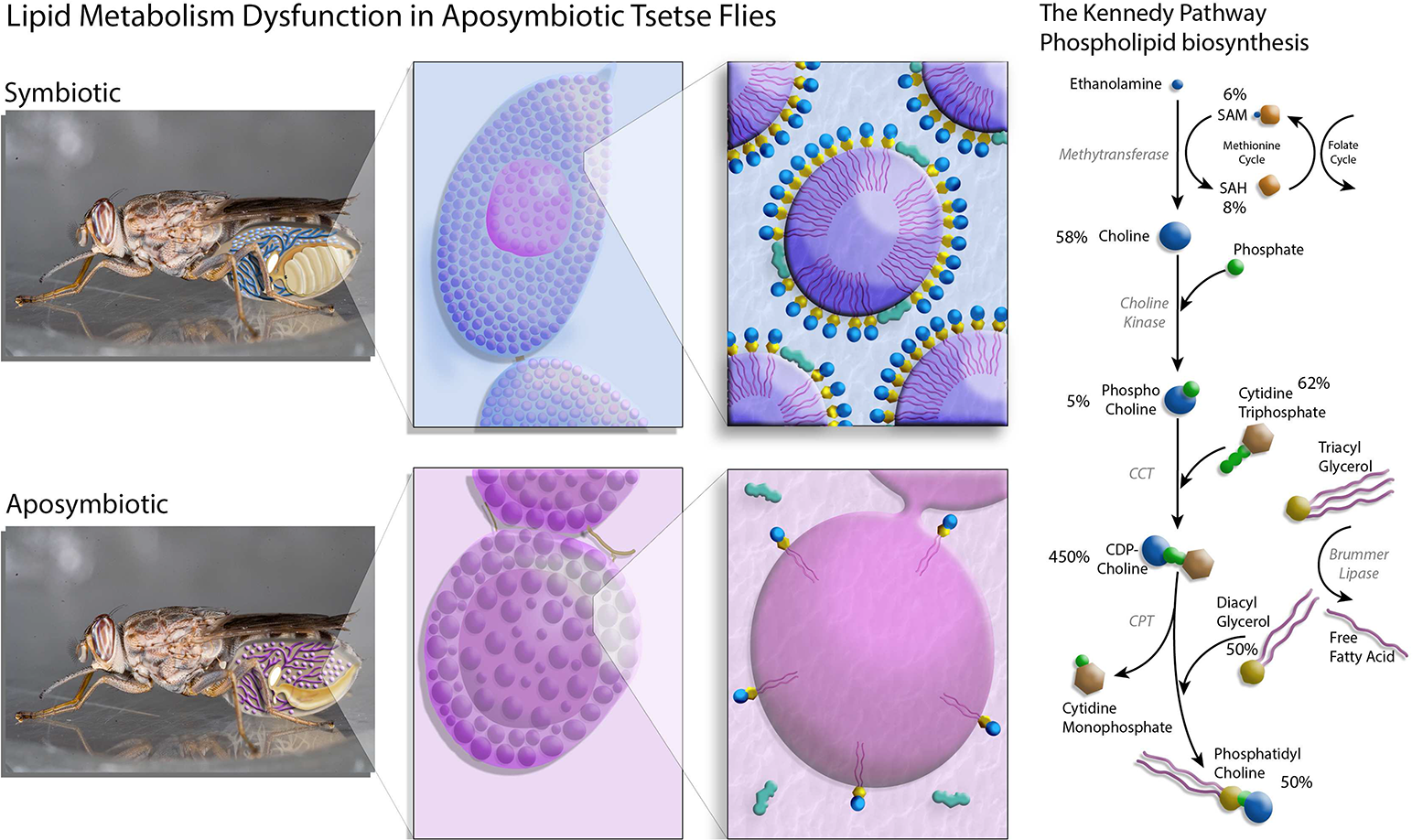
Summary of the dynamics between symbiont loss and reproduction during tsetse fly pregnancy in relation to altered lipolysis. The percentage annotations in the panel depicting the Kennedy Pathway represent the relative amount of the specific lipid metabolite in the aposymbiotic tsetse fly relative to the untreated, wild-type control which would be equivalent to 100%.

In mammals, disruption of appropriate PC to PE ratios in LDs causes formation of large droplets with reduced surface area and altered surface protein composition. The end result is abnormal fat retention within a cell or tissue/organ (steatosis; (Listenberger et al., 2018). In tsetse flies, symbiont clearance produces obesity, similar to that observed in mammals where individuals see increased lipid levels in conjunction with symbiont dysfunction. Flies initially fed blood spiked with tetracycline are cleared of their *Wigglesworthia*, with any lingering bacterial-derived factors clearing through subsequent feedings. Importantly, multiple feeding events following tetracycline treatment over the course of two weeks reduces the direct impact of this treatment, indicating that the phenotypes are associated with the loss of the symbiont. Symbiont absence results in PC deficiency that inhibits metabolism, mobilization and integration of stored lipids during lactation. Reproduction is terminated due to lipid retention in the FB and inefficient lipid transfer to the intrauterine larva. This is likely due to the combined effect of the mothers inability to process FB lipids due to dysfunction in lipid droplet formation in the FB and MG as *cct1* is abundantly expressed in both locations.

The biochemical observations associated with this phenotype suggest at least two points of failure in phosphatidylcholine biosynthesis via the Kennedy (CDP-choline) pathway. The first limiting factor appears to be deficiencies in choline and choline phosphate levels, which form the phosphatidylcholine headgroup. The production of choline is dependent on methylation reactions driven by enzymes that utilize S-adenosyl methionine (SAM) as a cofactor. Prior work in aposymbiotic *Glossina* demonstrates a severe deficiency of SAM in aposymbiotic flies (Bing et al., 2017). Production of SAM is dependent on the cysteine-methionine pathway, which is dependent on the B vitamins pyridoxal and folate as enzymatic cofactors (Loenen, 2006; Nguyen et al., 2001). Methylation reactions require SAM as a cofactor for the production of choline and conversion of phosphatidylethanolamine (PE) to PC (Gibellini and Smith, 2010; Mazen Noureddin et al., 2015; Snyder and Rio, 2015). In accordance with the observations in mammalian systems, the mechanistic disruption occurring in the aposymbiotic flies is most likely due to the deficiency in symbiont-derived B vitamins.

*Wigglesworthia* produces an array of B vitamins (B1, B2, B3, B5, B6, B7 and B9) (Akman et al., 2002; Bing et al., 2017; Rio et al., 2019; Snyder and Rio, 2015). Deficiencies in B6, folate and, consequently, SAM have a negative impact on phospholipid biosynthesis in mammalian systems. Issues related to the production of phosphatidylcholine have been linked to folate restriction (Abratte et al., 2008). The aposymbiotic tsetse phenotype bears resemblance to the pathology observed in non-alcoholic fatty liver disease (NAFLD) in mammals as PC deficiency is a key dysfunction (Bianca M. Arendt et al., 2013; Jacobs et al., 2013; Ville Männistö et al., 2019). Rats exhibiting a NAFLD phenotype have been rescued by dietary supplementation with phosphatidylcholine (Kitagawa et al., 2015). The phenotype observed in aposymbiotic flies (this study and in prior studies (Bing et al., 2017; Snyder and Rio, 2015)) is phenotypically equivalent to that observed when *cct1* is suppressed. Folate deficiency is likely a major underlying factor for the phenotype of dysfunctional metabolism of lipids we observe in flies lacking symbionts. Induction of folate deficiency in tsetse through treatment with glyphosate impacts vector competency and reproduction (Rio et al., 2019; Snyder and Rio, 2015), highlighting that folate provision by *Wigglesworthia* represents a critical lynchpin in tsetse metabolism. While female tsetse are able to survive B vitamin deficiency, the metabolic deficiencies prevent them from performing the mass mobilization of lipids from FB stores to the MG required for milk production during pregnancy. Our supplementation of B vitamins to aposymbiotic flies reduces obesity and has been previously shown to recover fecundity (Michalkova et al., 2014b), providing a direct link between B vitamins provided by *Wigglesworthia* and obesity in *Glossina*. This is not surprising as *cct* is overexpressed in aposymbiotic lines, indicating once specific B vitamins are available lipolysis can occur more rapidly to support reproduction.

Another disruption in phosphatidylcholine biosynthesis appears to derive from DAG deficiency. The lack of DAGs results in inadequate substrate availability for fusion with cytidine 5’-diphosphocholine by DAG cholinephosphotransferase. DAGs can be synthesized *de novo* during the production of triglycerides or they can be produced during lipolysis of triglycerides upon removal of a fatty acid side chain by a lipase. Triglycerides are overabundant in aposymbiotic flies, suggesting that lipolysis of the existing triglycerides is compromised under these conditions. In insects, the lipase responsible for this activity is called *brummer lipase* (*Drosophila*, CG5295), and is orthologous to the Adipose triglyceride lipase (ATGL) in mammals (Alexander Mildner et al., 2005; Grönke et al., 2007). Brummer and ATGL are both regulated by the phosphorylation state of other lipid droplet bound proteins, called perilipins (Sztalryd and Brasaemle, 2017). Perilipins sequester a co-activator of ATGL in their non-phosphorylated state called CGI-58, which inhibits lipolysis (Itabe et al., 2017). Recent work demonstrates that LDs lacking proper phosphatidylcholine levels show increased levels of associated perilipins and a steatosis (lipid accumulation) phenotype (Listenberger et al., 2018). The phosphatidylcholine deficiencies in aposymbiotic *Glossina* could be functioning via the same paradigm, resulting in perilipin accumulation and inhibition of *brummer*-mediated lipolysis resulting in DAG deficiency and TAG accumulation.

The symbiosis between tsetse flies and *Wigglesworthia* is essential for this complex system and both are dependent on one another for their survival. These studies demonstrate that lipid metabolism, a foundational process in *Glossina* biology, is impaired in the absence of their obligate symbionts. This is likely a model for other symbiotic systems, particularly those compensating for dietary nutritional deficiencies such as blood feeding invertebrates that are supplied with B vitamins from bacterial symbionts. This includes ticks (Duron et al., 2018), bed bugs (Nikoh et al., 2014), and lice (Boyd et al., 2017), which are all epidemiologically important blood-feeding disease vectors and require obligate symbionts to provide B vitamins. Beyond blood-feeding arthropods, specific bacteria associated with plant feeding insects also provide B vitamins (Salem et al., 2014), suggesting that lipid metabolism and B vitamin dependency issues likely occur in many systems. Lastly, B vitamins can be provided to insect species that have transient bacterial symbionts, such as in mosquitoes (Wang et al., 2021), suggesting that the interdependence of B vitamins and lipid metabolism may be a widespread phenomenon among arthropod species. As a key example, mosquitoes display altered lipid metabolism related to microbiome elimination and B vitamin deficiency (Didion et al., 2021; Romoli et al., 2021; Valzania et al., 2018; Wang et al., 2021). Specifically, the mosquitoes fail to accumulate lipids at critical time points, such as early in development and when lipids are accumulated for dormancy (Didion et al., 2021; Romoli et al., 2021; Valzania et al., 2018; Wang et al., 2021), indicating a critical role for B vitamins, lipid metabolism, and symbiont presence for many biological processes, including the critical contribution to reproduction we observed in this study.

Along with the role among arthropod systems, genomic studies have identified significant functional convergence among live-bearing systems in invertebrate and vertebrate lineages (Fouks et al., 2022). This includes aspects that range from metabolism to immune function (Fouks et al., 2022). In the vertebrate systems, fetal development requires essential fatty acids and long chain polyunsaturated fatty acids (Herrera, 2002; Knopp et al., 1998; Ryckman et al., 2015). As in the insect system, the accumulation of lipids early in vertebrate pregnancy is a critical feature that builds up stores providing lipids to the developing progeny (Herrera, 2002). As pregnancy progresses, the mother transitions from lipogenesis to lipolysis to increase the lipid supply to the fetus (Herrera et al., 1994, 1988). An imbalance in lipid levels provided to the neonates, either through malnutrition of the mother or impaired transfers, can yield many negative consequences ranging from delayed growth/development to death (Furness et al., 2012; Herrera, 2002). Similar to this study, B vitamin levels have been documented to alter lipid metabolism in mammalian systems with most focus on human and murine systems (Castaño-Moreno et al., 2020; da Silva et al., 2020; Herrera, 2002). Low B vitamin levels during pregnancy have been directly associated with a reduction in circulating lipid levels, increased obesity in the mother, and a reduction in health of the neonate (Adaikalakoteswari et al., 2017; Castaño-Moreno et al., 2020; McNeil et al., 2009), which is functionally similar to the obesity and reduced larval growth we observed in tsetse flies lacking B vitamins produced by *Wigglesworthia*. As observed in this study for tsetse flies, the induction of obesity acts through impaired PC synthesis that requires specific B vitamins, which is also critical for vertebrate lipid metabolism (da Silva et al., 2014). Thus, a distinct commonality of viviparity among animals is the requirements of specific B vitamins for lipid metabolism to meet the energetic demand of internal progeny development. This expands on genomic similarities (Fouks et al. 2022) to other physiological processes that are similar between invertebrate and vertebrate viviparity.

## Methods

### Flies

*Glossina morsitans morsitans* flies were maintained on defibrinated bovine blood (Lampire Biologicals, Pipersville, PA, USA) at 25°C and 50-60% RH using an artificial membrane feeding system (Moloo, 1971) at the insectaries at Yale School of Public Health and the Slovak Academy of Sciences. Females were mated 3-5 days after emergence. Flies were collected according to established developmental markers based on oocyte, embryo and larva presence (Attardo et al., 2006; Strickler-Dinglasan et al., 2006; Yang et al., 2010). Aposymbiotic tsetse larvae were derived from females fed a diet supplemented with tetracycline (20 μg per ml of blood) to clear their indigenous microbiota and yeast extract (1%) to provide vitamins in the absence of obligate *Wigglesworthia* (Pais et al., 2008). A subset of the progeny were allowed to feed of blood supplemented with B vitamins and yeast extract according to Michalkova et al. (2014b), which has been shown to partially alleviate B vitamin deficiencies following *Wigglesworthia* elimination.

### Milk extraction

Milk products were extracted from the guts of developing 3rd instar larvae based on our previous studies (Benoit et al., 2014, 2012). Briefly, actively-feeding individuals were removed from the uterus and their digestive tracts were dissected. A pulled glass capillary needle was utilized to pierce into the digestive tract and the contents were removed using reverse pressure. Each sample was stored at −80°C until utilization in the experiments.

### Lipidomics

Tissues (milk glands and fat body) were dissected from aged-matched WT flies that received normal blood meals for five weeks (control - symbiotic) and aposymbiotic females and prepared according to our previously established protocol (Bing et al., 2017). Briefly, tissues were added 100 μl of ice cold 1X phosphate buffered saline (PBS), homogenized with a pestle and stored at −80° C. For both aposymbiotic and control flies, samples consisted of four replicates of 20 individuals each. Samples were shipped to Metabolon (Morrisville, NC, USA) and analysis was performed using the proprietary Metabolon DiscoveryHD4 global lipidomics platform. Each sample was screened for the presence and quantity of lipids by mass spectrometry using a panel specific lipids. Relative differences between samples were determined by comparison of mean metabolite abundance across the biological replicates. Significance of the difference between symbiotic and aposymbiotic samples was determined by Welch’s two-sample t-test: p-value cut off of (<0.05). False discovery rate (FDR) was tested for by q-value with a cut off of (<0.1). Non-Metric Multidimensional Scaling (NMDS) plots were created using the metaMDS function from the R package vegan (Dixon, 2003; Oksanen, 2007) to determine differences in the general lipid composition between control and aposymbiotic individuals. Analysis of similarities (ANOSIM, using the anosim from the R package vegan) tests was performed to test for significance in the general differences.

### RNA extraction and qPCR

RNA was extracted from flies with TRizol reagent (Invitrogen, Carlsbad, CA, USA) using the manufacturer’s recommended protocol. RNA was treated with TURBO DNA Free kit (Ambion, Austin, TX, USA). RNA was precipitated with alcohol to remove residual salt, purified with a RNeasy kit (Qiagen, Maryland, USA) and stored at −70°C until use. cDNA was prepared using the Superscript III reverse transcriptase (Invitrogen). Transcript abundance was determined utilizing quantitative PCR (qPCR) on a Bio-Rad CFX detection system (Hercules). Primer sequences used were those from Table S1. Results were analyzed with CFX manager software version 3.1 (Bio-Rad). Ct values for genes of interest were standardized by Ct values for the control gene (tubulin) and relative to the average value for the control treatment or newly emerged flies, yielding a relative fold change in gene expression.

### RNA interference of *cct1*

Suppression of *cct1* was performed utilizing double stranded RNA (dsRNA) according to our previous studies on tsetse gene suppression (Benoit et al., 2017, 2012). The T7 promoter sequence was added to the 5′ end of the primer sequences and PCR amplification conditions are described (Table S1). The PCR products were purified using QIAquick PCR purification kit (Qiagen, Valencia, CA) and cloned into pGEM T-Easy vector (Promega, Madison, WI) and verified by sequencing (Keck DNA sequencing facility, Yale University). dsRNAs were synthesized using the MEGAscript RNAi Kit (Ambion, Austin, TX), purified using a RNeasy Mini Kit (Qiagen, Valencia, CA) and siRNAs were generated by using the Block-iT Dicer RNAi kit (Invitrogen, Carlsbad, CA). The siRNA concentration was adjusted to 600–800 ng/μl in PBS and each fly was injected with 1.5 μl siRNA using a pulled glass capillary needle into their thorax. Concentration of siRNAs were kept at 600–800 ng/μl within the 1.5 μl injected. Knockdown efficiency was determined 5d after injection by qPCR and normalized to *tubulin* levels. Fecundity following *cct1* was assessed according to previously established protocols (Benoit et al., 2014, 2012).

### Glyphosate treatment

Groups of *G. morsitans* flies were maintained on a diet containing 100 μM glyphosate 100 μM glyphosate combined with 500 nM folic acid [N-(phosphonomethyl)glycine; Sigma-Aldrich, St. Louis, MO, USA] or (Sigma-Aldrich). This treatment has been previously documented to impact folate synthesis (Rio et al., 2019; Snyder and Rio, 2015). Following six bloodmeals, the flies were examined to determine if lipid and phosphotydilcholine levels were increased or decreased. These treatments were conducted to eliminate off-target effects of tetracycline supplementation validate Wigglesworthia’s role in tsetse lipid metabolism provide a secondary validation of impaired lipid metabolism without treatment with antibiotics.

### Total lipid assay

Total lipids present within biological samples were determined with a standard vanillin assay (Attardo et al., 2012; Rosendale et al., 2019; Van Handel, 1985). Whole flies and individual tissues were dried at 0% RH at 60°C and weighed to determine the dry mass. The flies were homogenized in 2 ml of chloroform:methanol (2:1). The supernatant was removed and placed into a glass tube and the solvent was evaporated at 90°C. The dried lipids were treated with 0.4 ml of concentrated sulfuric acid at 90°C for 10 min. The acid/lipid mixture (40 μl) was combined with 4 ml vanillin reagent. Samples were measured spectrophotometrically at 525 nm and the total lipid content was calculated against a lipid standard.

### *In situ* hybridization for *cct1*

Milk gland tubules and fat body were removed from tsetse fly females at six-seven days after the previous birth. The combined milk gland/fat body was placed into Carnoy’s fixative for a five-six day fixation period (Attardo et al., 2008; Michalkova et al., 2014a). Antisense/sense digoxigenin-labeled RNA probes for *cct1* were generated using the MAXIscript T7 transcription kit following manufacturer’s protocol (Ambion, Austin, TX, USA) using a primer set with a T7 primer (Table S1). Antibody solutions were made using α-Digoxigenin-rhodamine, Fab fragments (Roche Applied Science, Penzberg, Germany) for FISH probe detection (1:200 dilution) and rabbit α-GmmMGP (1:2500) antibodies. Alexa Fluor 488 goat α-rabbit IgG (Invitrogen) at a dilution of 1:500 was added as a secondary antibody for immunohistochemistry (Attardo et al., 2008; Michalkova et al., 2014a). Slides were mounted in VECTASHIELD Mounting Medium with DAPI (Vector laboratories Inc. Burlingame, CA, USA). Samples were observed using a Zeiss Axioskop2 microscope (Zeiss, Thornwood, NY, USA) equipped with a fluorescent filter and viewed and imaged at 400x magnification. Images were captured using an Infinity1 USB 2.0 camera and software (Lumenera Corporation, Ottawa, Ontario, Canada) and merged in Adobe Photoshop.

### Phosphatidylcholine assay

Phosphatidylcholine levels were assessed through the use of a colorimetric assay (Caymen Chemicals, Ann Arbor, MI, USA). Extracted milk from the guts of 3rd instar larvae (10 μl) was placed within the sample well of a 96 well plate. The colorimetric reaction was initiated by adding 100 μl of the reaction mixture. Following mixing for a few seconds, the plate was incubated at 37°C for 60 min and concentration determined by comparison to a phosphatidylcholine standard.

### Nile Blue staining

Lipid composition of fat body cells from control and aposymbiotic *Glossina* was analyzed microscopically using the lipophilic Nile Blue stain. Nile Blue is a differential stain, which results in charged phospholipids (fatty acids, chromolipids and phospholipids) being stained blue and neutral lipids (triglycerides, cholesterol esters, steroids) being stained red/pink. The method used here is a modified version of the protocol described by Canete and colleagues (Canete et al., 1983). Fat body tissues were dissected from flies reared as described in the Lipidomics section above. Dissected tissues were immediately transferred to ice cold 4% paraformaldehyde in PBS and allowed to fix overnight at 4°C. Fixed tissues were then rinsed in PBS followed by immersion in 0.1 mg/ml aqueous Nile Blue sulfate (Sigma-Aldrich) for 30 min. After staining, tissues were rinsed three times with PBS for 1 min per rinse to remove excess stain. Tissues were then mounted in glycerol on single depression concave microscope slides. Stained fat bodies were visualized using a Zeiss Axio Vert.A1 scope and images captured using the Zeiss Zen imaging software. Color differences were quantified by measuring the average amount of red and blue color in five 100 pixel squares within the center of each lipid droplet with the use of Adobe Photoshop.

### Statistical analyses

Lipidomics data were analyzed using the Metaboanalyst software package (https://www.metaboanalyst.ca/) (Pang et al., 2021). Metabolite abundance values represent peak area counts measured from compound peaks. Samples were normalized across samples using the sum of raw area counts for each sample. Missing values were imputed by replacement with minimum detected values for each metabolite. Data were rescaled by median centering to a value of 1 for each sample. Significant differences between control and aposymbiotic samples were measured by t-test followed by p-value correction using false discovery rate (FDR)/Benjamini and Hochberg analysis. Differences in relative abundance were considered significant below an FDR corrected P-value of 0.05. Differences in other biological attributes were assessed through the use of t-test or ANOVA.

## Funding

This study was supported by awards from National Institutes of Health USA (AI109263, AI128523, F32AI093023, and AI08177405), FAO/IAEA Coordinated Research Program “Improving SIT For Tsetse Flies through Research on their Symbionts” and “Improvement of Colony Management in Insect Mass-rearing for SIT Applications”, and the Cariplo-Regione Lombardia research grant IMPROVE (F.S.).

---

**Supplemental Figure 1:** Absolute abundance of detected fat body/milk gland phospholipids and associated compounds in symbiotic and aposymbiotic *G. morsitans.* Abundance values represent normalized area under compound peaks. Significance determined by Welch’s Two-Sample t-test followed by FDR adjustment of resulting p-values.

**Supplemental Figure 2:** Absolute abundance of detected fat body/milk gland diacylglycerols in symbiotic and aposymbiotic *G. morsitans.* Abundance values represent normalized area under compound peaks. Significance determined by Welch’s Two-Sample t-test followed by FDR adjustment of resulting p-values.

**Supplemental Figure 3:** Absolute abundance of detected fat body/milk gland sphingolipids and associated compounds in symbiotic and aposymbiotic *G. morsitans.* Abundance values represent normalized area under compound peaks. Significance determined by Welch’s Two-Sample t-test followed by FDR adjustment of resulting p-values.

**Supplemental Figure 4:** Absolute abundance of detected fat body/milk gland polyunsaturated fatty acids (PUFAs) in symbiotic and aposymbiotic *G. morsitans.* Abundance values represent normalized area under compound peaks. Significance determined by Welch’s Two-Sample t-test followed by FDR adjustment of resulting p-values.

**Supplemental Figure 5:** Absolute abundance of detected fat body/milk gland lysolipids in symbiotic and aposymbiotic *G. morsitans.* Abundance values represent normalized area under compound peaks. Significance determined by Welch’s Two-Sample t-test followed by FDR adjustment of resulting p-values.

**Supplemental Figure 6:** Absolute abundance of detected fat body/milk gland amino acid and vitamin B associated metabolic pathways undergoing significant changes in abundance between in symbiotic and aposymbiotic *G. morsitans.* Abundance values represent normalized area under compound peaks. Significance determined by Welch’s Two-Sample t-test followed by FDR adjustment of resulting p-values.

**Supplemental Figure 7:** NMDS analysis comparing the lipid profile differences between aposymbiotic and control flies. Significant variation between aposymbiotic and control samples (ANOSIM, R = 0.879, P = 1.03E-06).

**Table S1:** Primers used for qPCR, generation of dsRNA, and *in situ* hybridization.

## Notes

### Competing Interest Statement

The authors have declared no competing interest.

### Summary of Updates

We have added a new portion of Figure 4.

