## Supplementary figures and images for "Lipid metabolism dysfunction following symbiont elimination is linked to altered Kennedy pathway homeostasis"

### Supplemental Figure 1

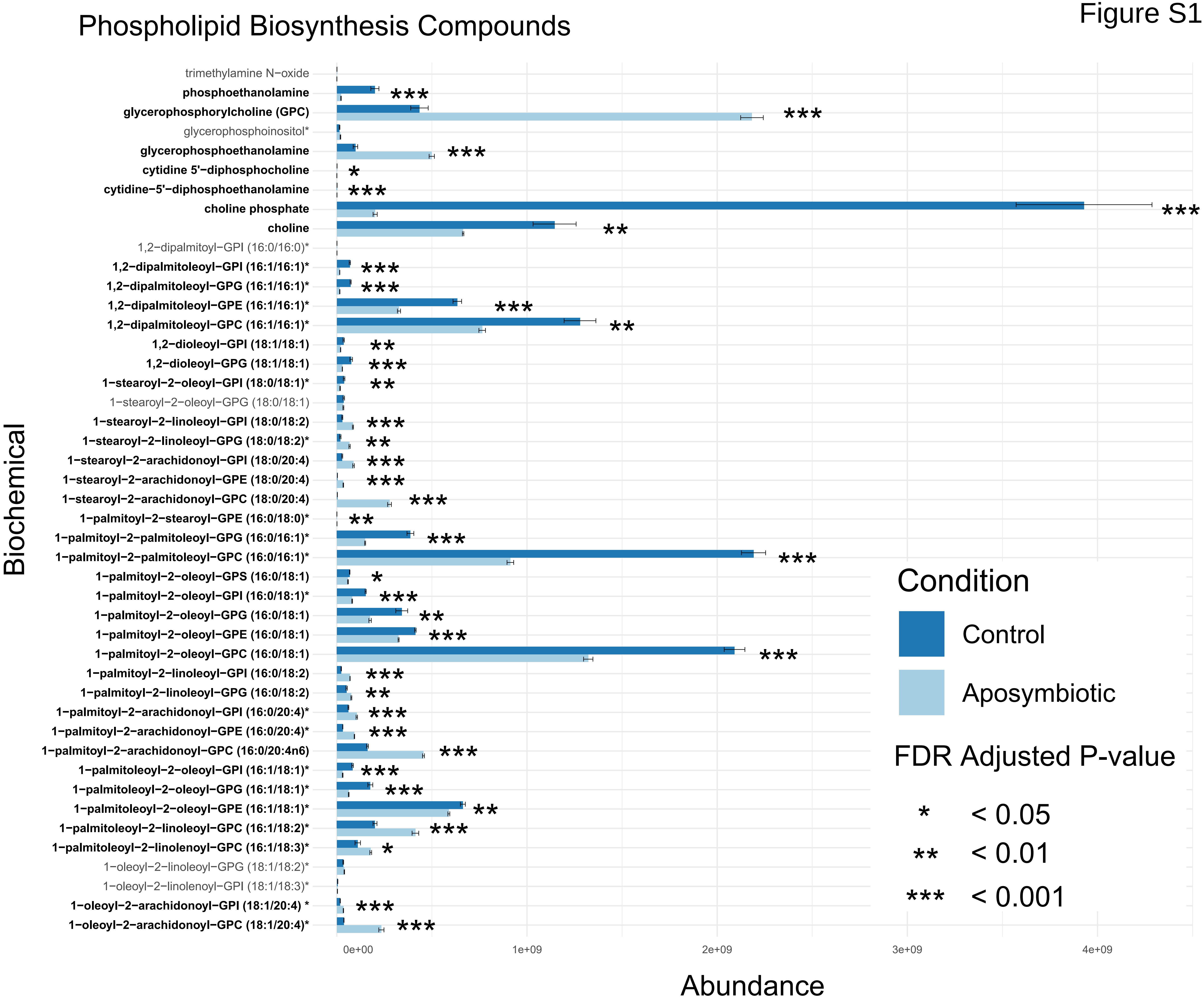

### Supplemental Figure 2

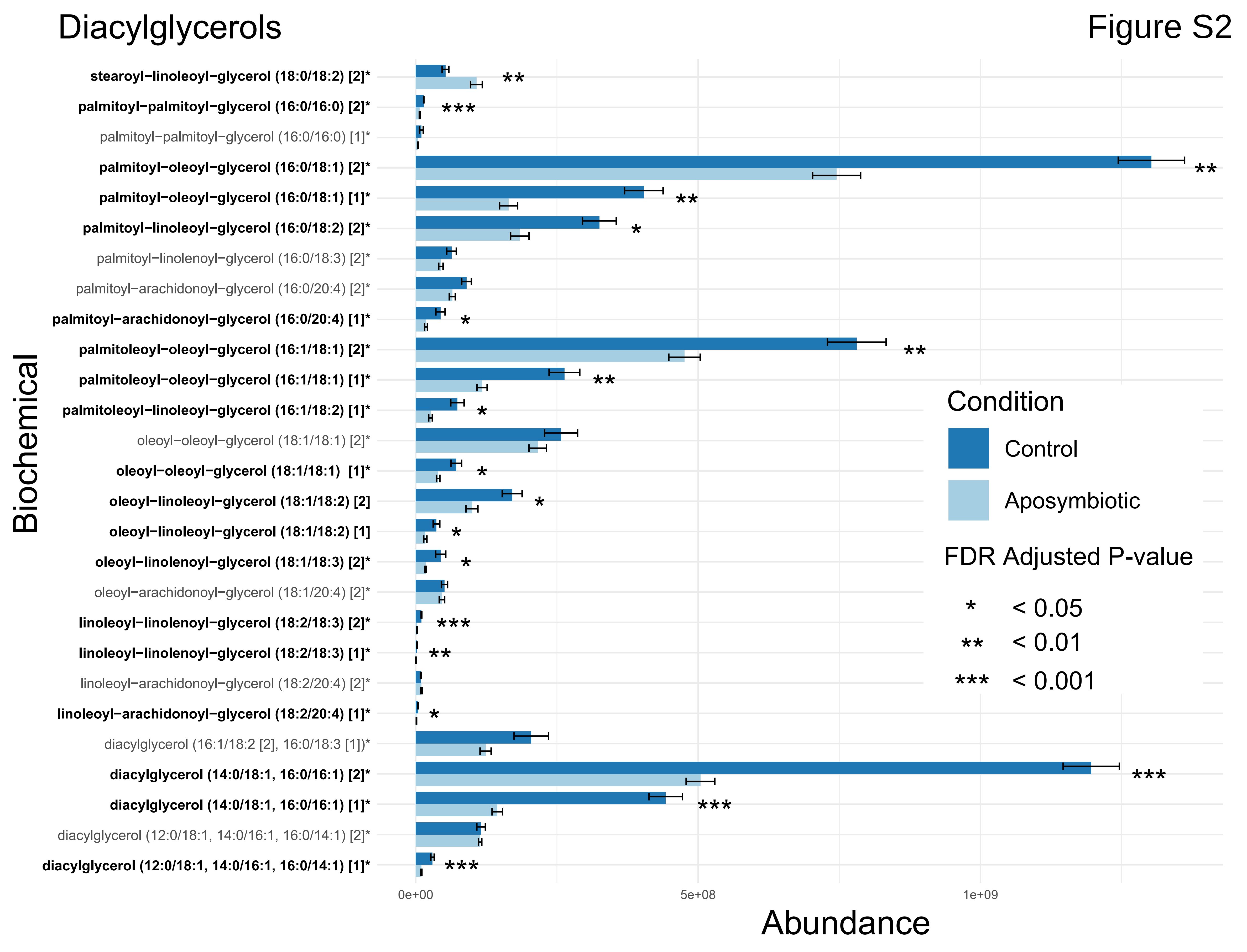

### Supplemental Figure 3

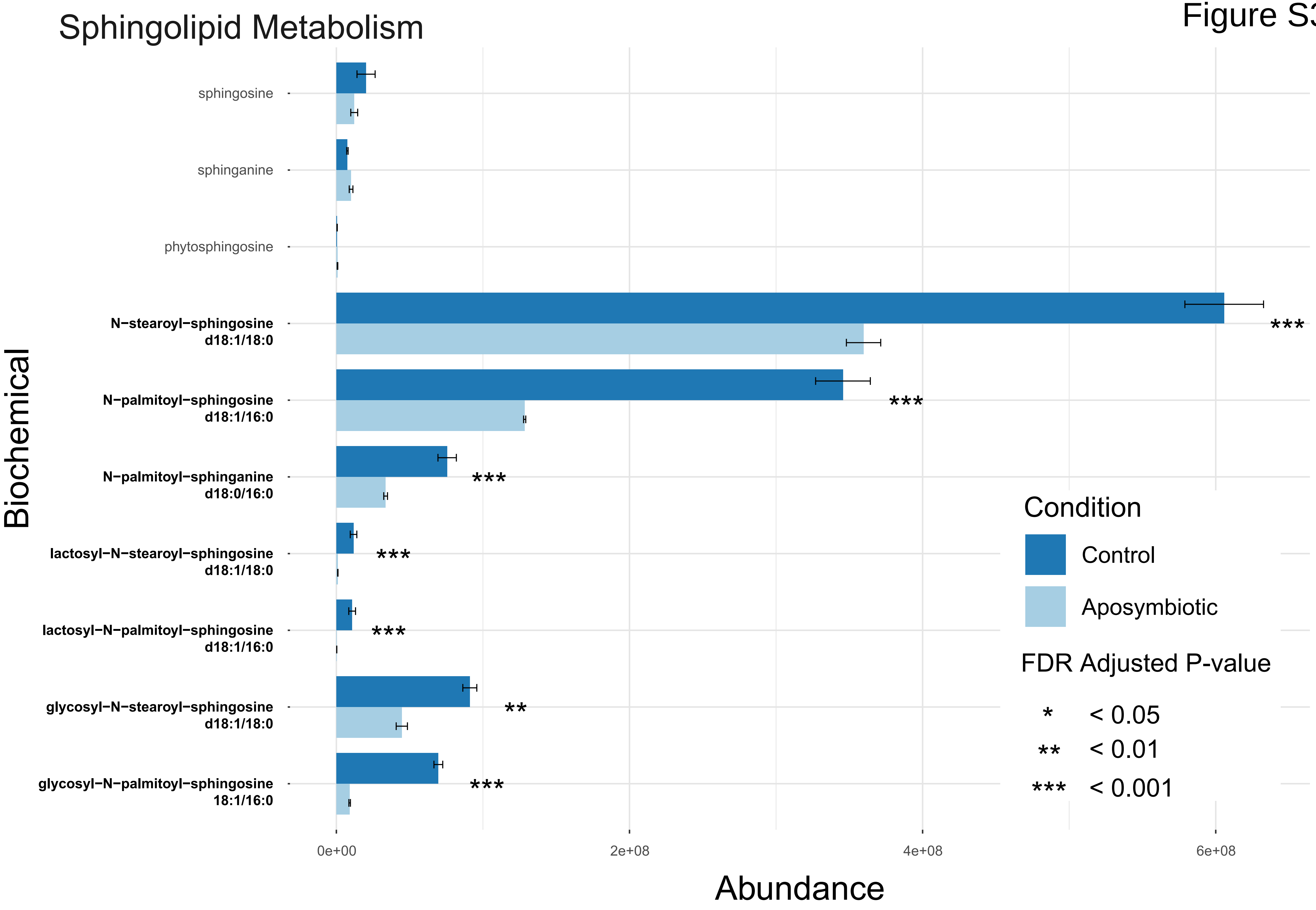

### Supplemental Figure 4

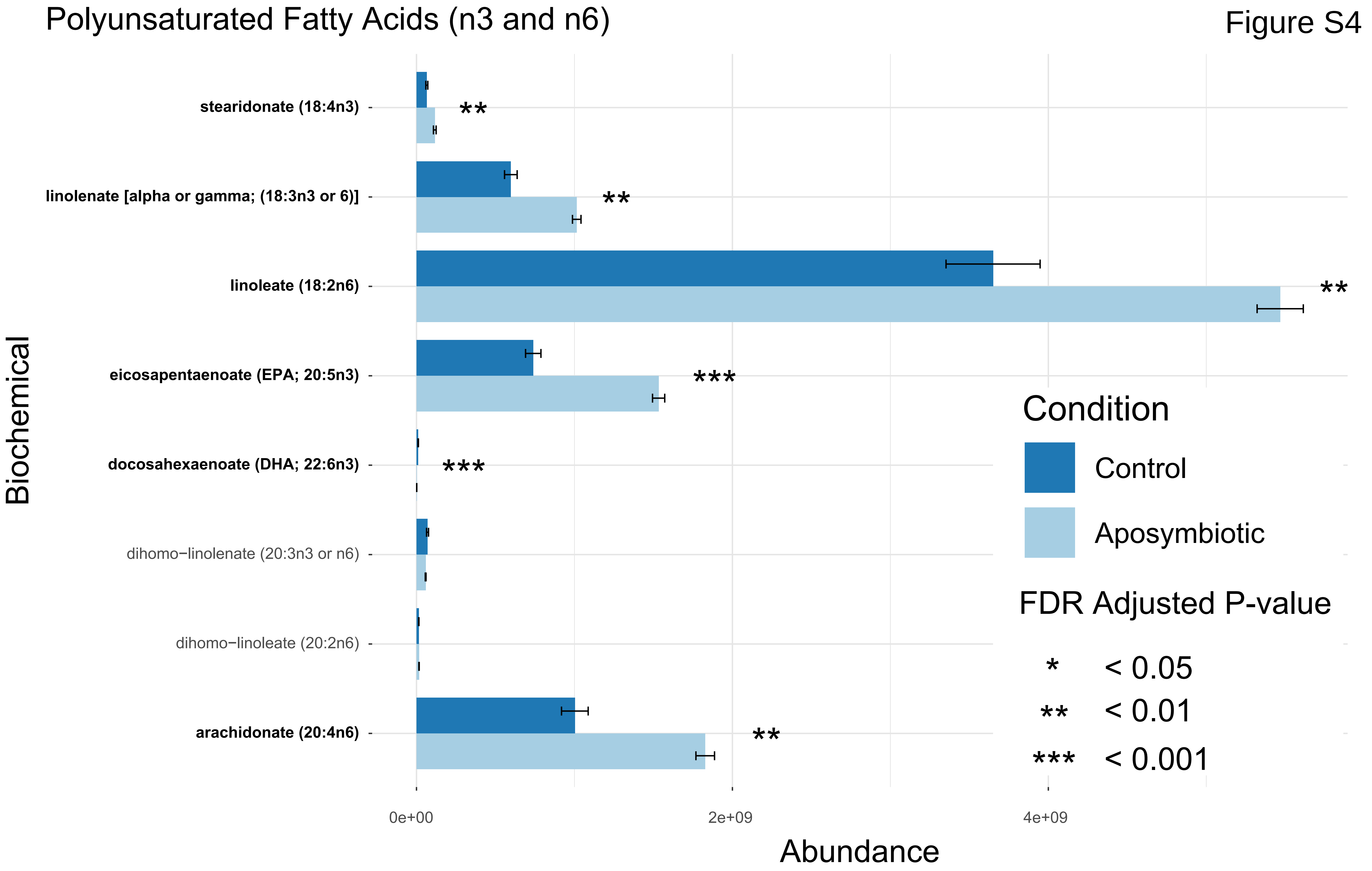

### Supplemental Figure 5

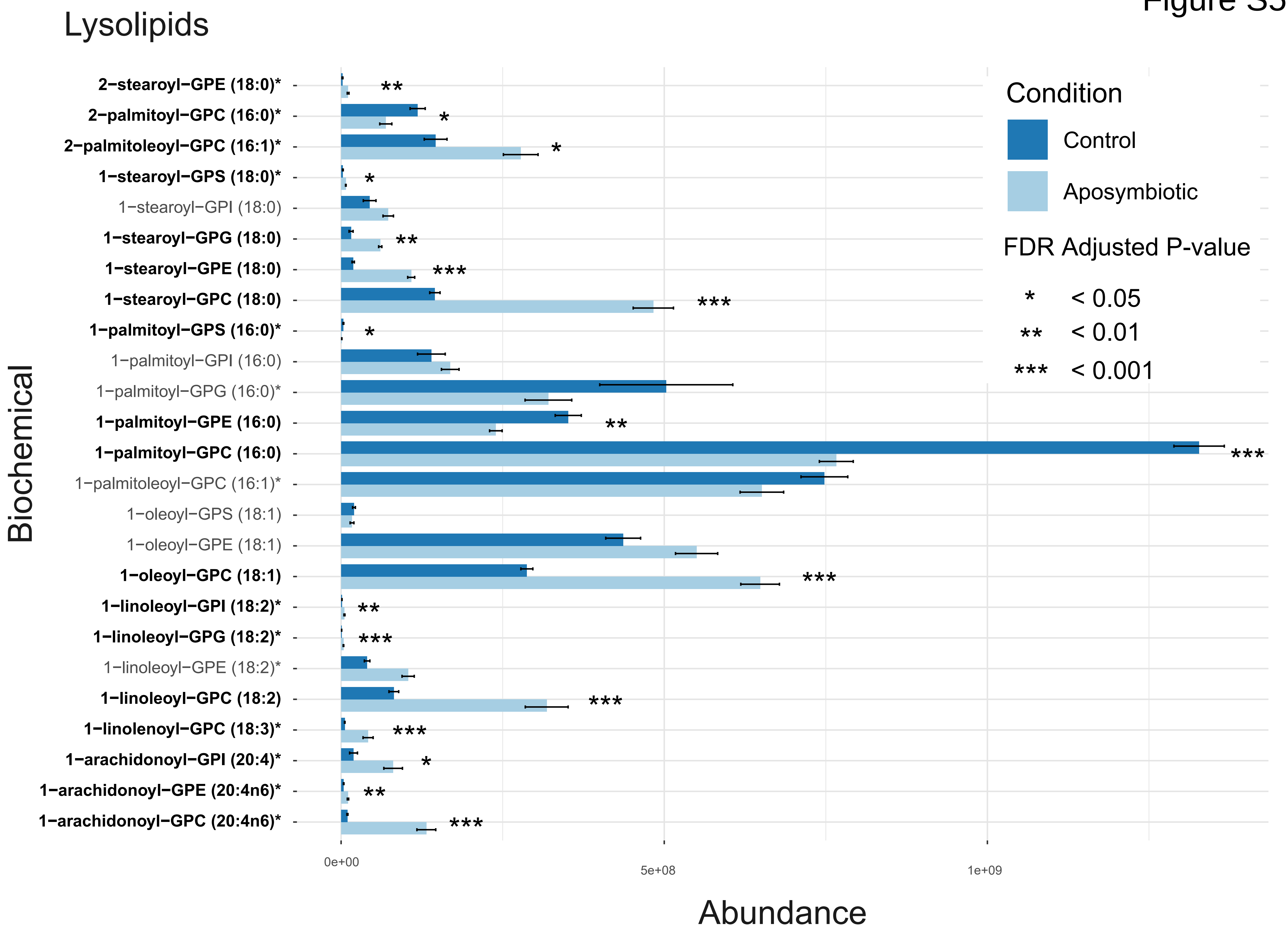

### Supplemental Figure 6

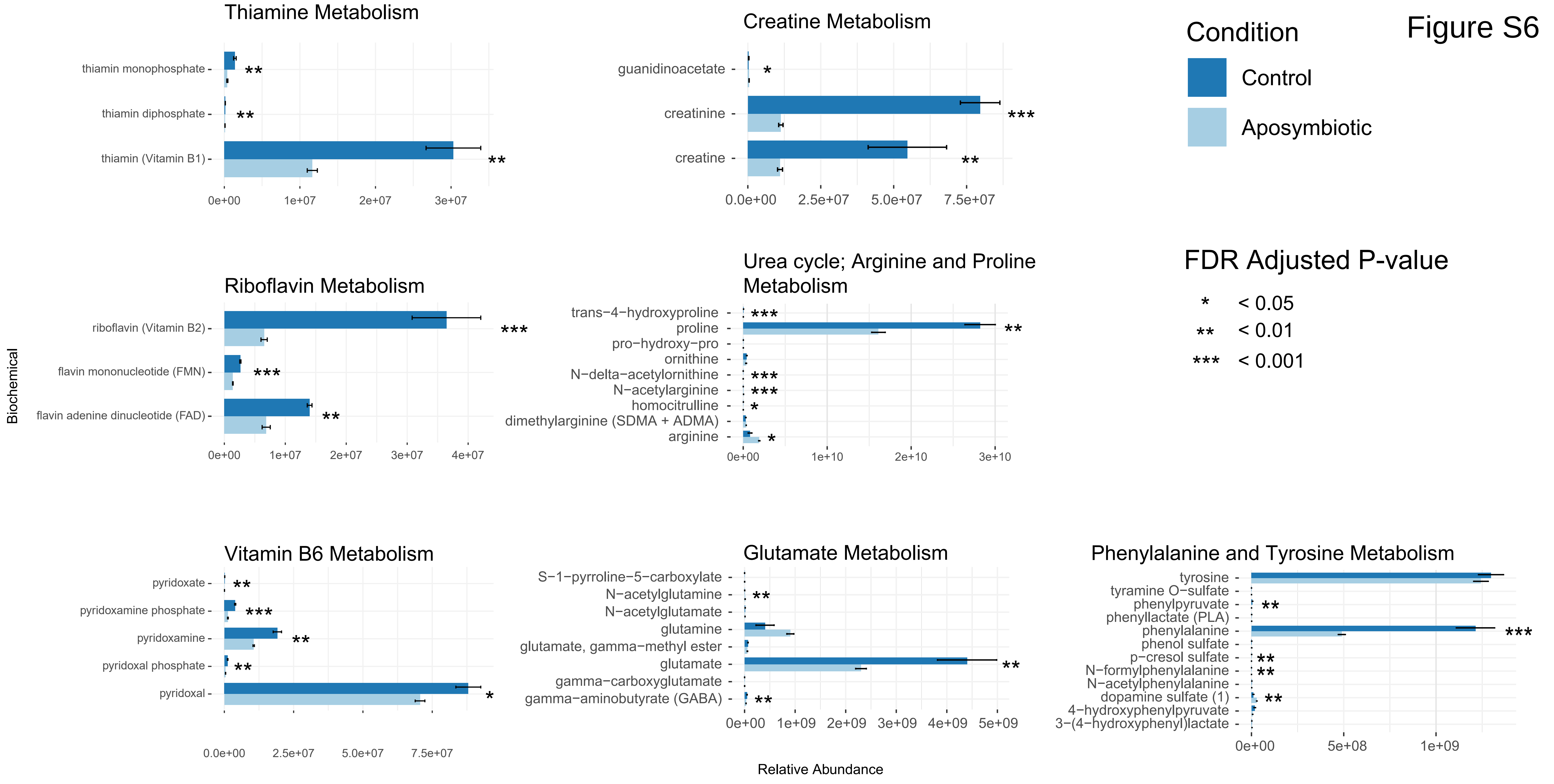

### Supplemental Figure 7

#### Figure S7

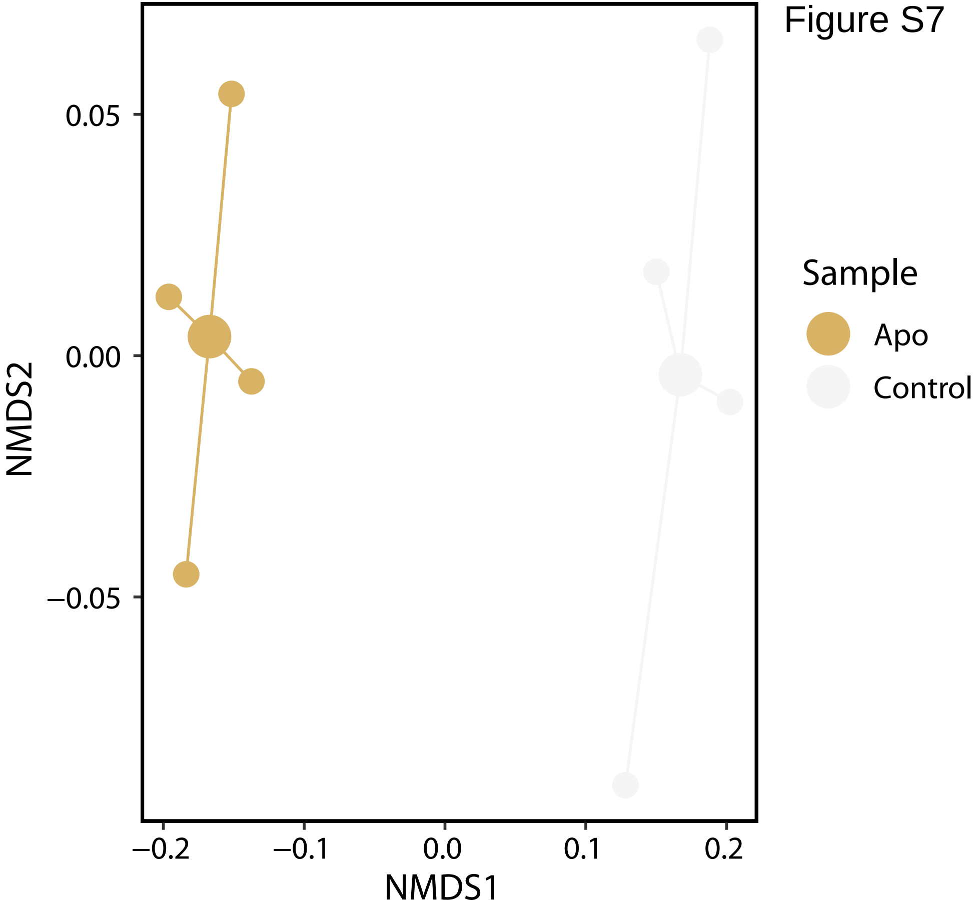
